# Exploring patterns of ongoing thought under naturalistic and task-based conditions

**DOI:** 10.1101/2020.07.29.226431

**Authors:** Delali Konu, Brontë Mckeown, Adam Turnbull, Nerissa Siu Ping Ho, Tamara Vanderwal, Cade McCall, Steven P. Tipper, Elizabeth Jefferies, Jonathan Smallwood

## Abstract

Previous research suggests that patterns of ongoing thought are heterogeneous, varying across situations and individuals. The current study investigated the influence of a wide range of tasks and individual affective style on ongoing patterns of thought. In total, we used 9 different tasks and measured ongoing thought using multidimensional experience sampling. A Principle Component Analysis of the experience sampling data revealed four patterns of ongoing thought. Linear Mixed Modelling was used to examine the contextual distribution of the thought patterns. Different thought patterns were found to relate to different types of conditions. Intrusive and negative thought pattern expression was found to be influenced by individual affective style (depression level). Overall, these data show that patterns of thought are subject to both contextual and intrinsic variation, suggesting that understanding these important features of experience across a broad range of situations will be useful in understanding their role in human experience.

**Highlights:**

- Patterns of thought vary across different task contexts
- Thought pattern expression is influenced by individual affective style
- There is a need to broaden the tasks used to study ongoing thought

## 1. Introduction

Patterns of ongoing experience are hypothesised to be influenced by both the environment and the intrinsic features of individuals, such as their cognitive expertise or their affective style. For example, studies show that complex task environments reduce the self-generation of personally relevant information, and increase patterns of cognition with detailed task focus (Turnbull, Wang, Murphy, et al., 2019). In addition, reading interesting texts help individuals maintain attention on the narrative, while texts that are more difficult show the opposite pattern (Giambra & Grodsky, 1989; Smallwood, Nind, & O’Connor, 2009; Unsworth & McMillan, 2013). Notably, however, recent work has demonstrated that patterns of ongoing thought in the context of the real-world have both similarities and differences with those observed in the laboratory (Ho et al., 2020; Linz, Pauly, Smallwood, & Engert, 2019). The disparity between patterns of thought in the lab and in the real world suggest that the types of tasks individuals often engage with in daily life may not correspond to those that are often used in experimental contexts. This may be particularly true for tasks like the sustained attention to response task; SART (Robertson, Ridgeway, Greenfield, & Parr, 1997), which engenders situations that maximise the need to maintain attention on task-relevant material with little or no support from the external environment. Paradigms such as the SART may provide a useful tool with which to study sustained attention, but they may not relate well to many of the everyday situations in which people generally spend their time.

Studies examining the role of intrinsic influences on patterns of ongoing thought highlight the relevance of individual differences in affective style and cognitive expertise. For example, individuals who are anxious or unhappy engage in greater off-task thought, often with repetitive or unpleasant features (Makovac et al., 2018; Ottaviani & Couyoumdjian, 2013). In the cognitive domain, individuals with a high capacity for executive control maintain attention more effectively during complex task environments (McVay & Kane, 2009; Unsworth & McMillan, 2013), and refrain from generating off-task thoughts until task environments are less demanding (Rummel & Boywitt, 2014; Smallwood et al., 2009; Turnbull, Wang, Murphy, et al., 2019). In contrast, individuals who excel at tasks that depend on memory tend to generate patterns of thought involving mental time travel with vivid detail (Wang et al., 2019). Individuals who do well on creativity tasks report high levels of daydreaming (Baird et al., 2012; Smeekens & Kane, 2016; Wang et al., 2018), and individuals with expertise in disciplines such as poetry or physics often identify solutions to problems when their mind wanders from the task they are performing (Gable, Hopper, & Schooler, 2019).

Together contemporary research highlights that ongoing experience is subject to influences that are related to both internal features of the individual and to external features of the task environment. Despite recognition that different environments or testing conditions may evoke differences in the content and pattern of ongoing thought, studies often examine the experience in the laboratory in only a small set of task environments. In the current study, we aimed to bridge this gap in the literature by examining how reported patterns of thought vary across a wide range of different task environments. We chose a range of conditions, ranging from simple paradigms that isolate a common cognitive process, to complex higher order tasks that rely on multiple task components (such as gambling or set-switching), and finally to more engaging, dynamic TV-based naturalistic conditions such as TV-based that conditions which more closely mimic the complexity of daily life (Vanderwal et al., 2017). To see whether thought reported during these tasks related to individual affective style we also measured levels of anxiety (state and trait) and depression in our participants since these have been linked to differences in both self-reported and psychophysiological correlates of thought patterns gained via experience sampling (Deng et al., 2012; Hoffmann, Banzhaf, Kanske, Bermpohl, & Singer, 2016; Makovac et al., 2018; Ottaviani et al., 2014; Poerio, Totterdell, & Miles, 2013; Smallwood, O’Connor, Sudbery, & Obonsawin, 2007; Xu, Purdon, Seli, & Smilek, 2017).

In our study, we used multidimensional experience sampling (MDES) to identify different features of thought patterns, using a set of questions that we used in a prior brain imaging study (Konu et al., 2020). In our prior study we examined how the different patterns of thoughts were associated with ongoing neural activity during a low demand sustained attention task using functional magnetic resonance imaging (fMRI). We found that reports of ongoing thoughts with episodic and social features were associated with increasing activity in a region of the ventral medial prefrontal cortex. In that study and others, in order to identify the patterns of thoughts, we used a Principal Components Analysis (PCA) to create a common low dimensional representation of the experience sampling data, thereby identifying “patterns of thought” (Konishi, Brown, Battaglini, & Smallwood, 2017; Turnbull, Wang, Murphy, et al., 2019; Vatansever, Bozhilova, Asherson, & Smallwood, 2019). In the current study, we use PCA to determine the dimensions that make up the matrix of our experience sampling reports and, use these as a guide to explore (i) how our tasks evoke different patterns of thought and (ii) whether any of these patterns are also related to measures of the individuals’ affective style assessed via questionnaire. To understand how the task environment influences thought patterns, we used a Linear Mixed Model (LMM) in which the patterns of thought were compared across the different task environments. To understand the impact of individual variation on experience we performed a Repeated Measures Analysis of Variance (ANOVA) to examine whether the distribution of the thought patterns were associated with our participants’ affective style (anxiety and depression).

## 2. Methods

### 2.1. Participants

Seventy participants took part in a two-part behavioural study (60 females; mean age: 20.60 years; standard deviation: 2.10 years, age range: 18-34 years). All participants were native English speakers with normal/corrected vision between the ages of 18 and 35. This cohort was acquired from the undergraduate and postgraduate student population at the University of York. The study was approved by the local ethics committee at the University of York’s Psychology Department. All volunteers provided informed written consent and received monetary compensation or course credit for their participation.

### 2.2. Procedure Overview

The task paradigms reported in the current study were part of a larger cohort collection that tested the influence of situational and intrinsic influences on ongoing thought. The current study involved 4 hours of testing split over 2 separate sessions on consecutive days for ~2 hours each. Order of session and task was counterbalanced across participants. The order in which participants completed these tasks was pseudorandom using fixed order. Participants were tested in the same environment using the same computers and i-Pads over the two sessions.

In one session participants completed the TV-based paradigms (documentary and affective TV-based paradigms), multidimensional experience sampling and affective measure questionnaires. In the TV-based watching session, participants completed a passive documentary TV-based paradigm (3-4 minute TV-clips taken from the BBC TV program *Connections: Season 1*: see below for details) and a passive affective TV-based paradigm (3-4 minute TV-clips taken from BBC TV programmes *Happy Valley, Line of Duty, Luther and Bodyguard*: see below for details). In both TV-based watching paradigms participants were shown unique TV-clips and no clips were shown twice.

In a separate ‘task’ session, participants completed 7 tasks: the Go/No-go, Self, Semantic paradigms, as well as the Cambridge neuropsychological test automated battery (CANTAB® [Cognitive assessment software]. Cambridge Cognition (2019). All rights reserved. www.cantab.com) which consisted of the Cambridge Gambling, Emotional Recognition, Intra-Extra Dimensional Set-Shift and Spatial Working Memory task paradigms (see below for details). Participants also completed multidimensional experience sampling and affective measure questionnaires. A summary of the task paradigms used in the current study can be found in Table 1.

**Table 1.** Summary of task paradigms used in the current study with corresponding mean RT (ms), mean accuracy and standard error.

| Category | Task Paradigm | Condition | Task | RT (ms)<br>(standard error) | ACC<br>(standard error) |
| --- | --- | --- | --- | --- | --- |
| Simple tasks | Go/No-go | Go | Respond to nominated target | 435.27<br>(1.65) | 0.97<br>(0.01) |
|  |  | No-go | Respond to less frequent nominated target | 484.47<br>(2.80) | 0.96<br>(0.02) |
|  | Semantic | Visual Semantics** (Picture) | Make a decision (Europe or not) based on a pictorial stimulus | 808.32<br>(3.59) | 0.80<br>(0.01) |
|  |  | Verbal Semantics** (Word) | Make a decision (Europe or not) based on a text stimulus | 818.88<br>(3.64) | 0.81<br>(0.01) |
|  | Self-reference | Self reference | Make judgement in reference to self | 766.05<br>(3.14) | 0.91<br>(0.01) |
|  |  | Social cognition reference | Make judgement in reference to other | 774.49<br>(3.14) | 0.89<br>(0.01) |
| Complex tasks* | Working memory |  | Hold information in mind | N/A | N/A |
|  | Switching |  | Switch between different tasks | 81730.29<br>(3234.97) | N/A |
|  | Gambling |  | Make gambling decisions | 1325.955<br>(32.26) | N/A |
|  | Emotion Recognition |  | Identify emotional expressions | 1010.059<br>(25.51) | 0.66<br>(0.01) |
| TV-based tasks | Documentary TV-based clips | Documentary | Watch a documentary | N/A | N/A |
|  |  | Audiobook | Listen to a documentary | N/A | N/A |
|  |  | Audio Inscapes | Listen to documentary with irrelevant visual input | N/A | N/A |
|  | Affective TV-based clips | Suspense | Watch a TV clip in which a threat occurred at the end of the clip creating threat uncertainty | N/A | N/A |
|  |  | Action | Watch a TV clip in which the threat occurred at the start of the clip creating threat certainty | N/A | N/A |

\* From the CANTAB battery
\*\* From Rice, Hoffman, Binney, & Lambon Ralph, 2018

#### 2.2.3 Multidimensional Experiential Sampling (MDES)

Participant ongoing thought was measured using multidimensional experience sampling (MDES). Participants were asked 16 MDES questions as part of a larger experimental question, 13 of these questions have been retained for analysis in the current study. In the current study participants were asked how much their thoughts were focused on the task, followed by 12 randomly shuffled questions about their thoughts (Table 2). All questions were rated on a scale of 1 to 10. Within each block of all the tasks participants completed, one set of MDES probes yielding a total of 9 probes per individual for the documentary TV-based task, 8 probes per individual for the affective TV-based task, 12 probes per individual for the Go/No-go, Self and Semantic tasks, and 4 probes per individual for the CANTAB tasks, giving a total of 33 probes. Every block of each task was followed by MDES probes, probes were presented for 4 seconds maximum on the screen based on the average response time from previous studies, followed by a 500ms fixation cross. Participants were tested using different MDES sampling rates across the tasks due to difference in task block length.

**Table 2.** Multidimensional Experience Sampling questions used to sample thoughts in the current study.

| Dimension | Statements | Scale_low | Scale_high |
| --- | --- | --- | --- |
| Task | My thoughts were focused on the task: | Not at all | Completely |
| Future | My thoughts involved future events: | Not at all | Completely |
| Past | My thoughts involved past events: | Not at all | Completely |
| Self | My thoughts involved myself: | Not at all | Completely |
| Other | My thoughts involved other people: | Not at all | Completely |
| Emotion | The emotion of my thoughts was: | Negative | Positive |
| Modality | My thoughts were in the form of: | Images | Words |
| Detail | My thoughts were detailed and specific: | Not at all | Completely |
| Deliberate | My thoughts were: | Spontaneous | Deliberate |
| Problem | I was thinking about solutions to problems (or goals): | Not at all | Completely |
| Diverse | My thoughts were: | One topic | Many topics |
| Intrusive | My thoughts were intrusive: | Not at all | Completely |
| Source | My thoughts were linked to information from: | Environment | Memory |

*Participants rated statements from 1-10.*

#### 2.2.4 Affective measures

To gain an understanding of the individuals’ affective style, we administered the Centre for Epidemiological Studies Depression Scale; CES-D (Radloff, 1977) as well as the State and Trait Anxiety Inventory; STAI (Spielberger, 1983). The measures were completed at the end of the task session. These questionnaires were administered to participants using Qualtrics software (© 2019 Qualtrics). Qualtrics and all other Qualtrics product or service names are registered trademarks or trademarks of Qualtrics, Provo, UT, USA. https://www.qualtrics.com. The questionnaires used in the current study were collected as part of a larger battery of questionnaires designed to test differences in intrinsic influences on thoughts.

#### 2.2.5. Documentary TV-based paradigm

In the passive documentary TV-based paradigm, participants were instructed to attend to the screen as they watched and listened to TV-clips from a British documentary series (BBC TV program), called *Connections: Season 1* which reviews the history of science and innovation. Clips were presented under three audio-visual conditions: (i) congruent visual and auditory presentation (documentary condition) in which participants watched and listed to the documentary TV-clips, (ii) audio condition in which participants had audio input of the documentary clip accompanied by a white fixation cross, and (iii) *Inscapes* in which participants had audio input of the documentary clip with visuals from *Inscapes,* a nonverbal, non-social TV paradigm that features slowly moving abstract shapes from Vanderwal and colleagues (Vanderwal et al., 2017; Vanderwal, Kelly, Eilbott, Mayes, & Castellanos, 2015). *Inscapes* was shown to provide irrelevant yet complex dynamic visual input that was unrelated to the audio from the documentary. *Inscapes* was slowed to half speed and segmented into 3 unique clips so that the participants did not see the same clip twice. The order of audio-visual conditions was pseudo-randomised, so that 3 consecutive TV-clips always included one from each condition. Each session consisted of a total of 9 TV-clips. Participants were informed that they would watch documentary TV-clips with varying visual input but were unaware of which condition they were in before starting the block. Written instructions were presented at the start of each run. Participants were asked questions about the content of the TV-clips in a comprehension questionnaire at the end of the session (this data was part of the larger cohort collection and has not been used in current study). Seven participants were informed that they would be required to perform this questionnaire before the protocol was changed so that remaining participants were unaware that this was required.

#### 2.2.6. Affective TV-based paradigm

In the affective TV-based paradigm, participants were instructed to attend to the screen as they watched and listened to 3-minute TV clips from the BBC TV programmes *Happy Valley, Line of Duty, Luther and Bodyguard*, a range of commercial television shows including crime dramas and thrillers. The clips were intentionally chosen to each include a threatening event. There were two conditions which varied in the onset of the threatening event; i) an action condition in which the direct threat occurs in the first minute of the clip and the rest of the clip follows the protagonist(s) responses to the threat and ii) a suspense condition in which a potential threat, high in uncertainty is detected early on in the clip but the direct threat only occurs in the last minute of the clip, as discussed in McCall and Laycock (in review). Three independent raters were used to identify when the direct threat occurred in each clip. An example of an action condition clip is a scene from *Bodyguard Series 1 Episode 2* in which gunshots from a roof are fired (threatening event) at the protagonists within the first minute of the clip and the remainder of the clip follows the protagonists’ reaction to continuing shots. An example of a suspense condition clip is a scene from *Luther Series 3 Episode 2* in which two characters hear a noise when they believe they are home alone and go upstairs to investigate, in the last minute of the clip the characters are attacked (threatening event after a period of suspense). After each clip, participants were invited to (a) take a break for as long as they needed and (b) withdraw from the task if they were feeling distressed. Participants were asked to fill out a questionnaire on Qualtrics (Qualtrics, Provo, UT) regarding the TV clips watched and the state anxiety questionnaire from the state-trait inventory (STAI) just before and just after the affective TV-based paradigm. These questionnaires were part of the larger cohort collection and are not considered in the current study.

The order of the affective TV conditions was pseudo-randomised so that the first TV clip seen by participants was either from the action or suspense condition and this was counterbalanced across participants. The remaining TV clips were pseudo-randomised so that each condition would not be shown more than twice consecutively. Each session consisted of a total of 8 TV clips. Written instructions were presented at the start of each run. Participants were informed that the clips involved dangerous behaviour, strong language, and violence on several occasions prior to starting and they were reminded repeatedly that they had the right to withdraw at any time, without giving reason and without prejudice.

#### 2.2.7. PsychoPy tasks

PsychoPy3 (Peirce et al., 2019) was used to present the Go/No-go, Self and Semantic task paradigms to participants. Each task included 4 task blocks (2 blocks consisting of each experimental condition) and lasted ~3 minutes. Key press across all task paradigms was counterbalanced with participants making forced choice responses using d and k to indicate ‘yes’ or ‘no’ respectively. In the Semantic and Self tasks, a probe preceded stimulus presentation where each trial consisted of a probe signalling whether the trial was experimental or control. In each task paradigm trials consisted of the presentation of a target stimulus until a response was made (1500ms). Once a response was captured, a fixation cross appeared on the screen for the remaining time. The inter-stimulus-intervals (ISI) consisted of a fixation cross and was jittered (500-1500ms). Block order was counterbalanced across participants. Written instructions were presented at the start of each block.

##### 2.2.7.1 Go/No-go task paradigm

Participants were instructed to attend to the centre of the screen as a single shape stimulus was presented (‘X’, ‘Q’ or ‘O’). In the Go condition participants were instructed to make a single key press when the target stimulus ‘X’ was presented. In the No-Go condition the ‘O’ was the target stimulus to which participants had to make a single key press. Each block of the experiment was designed so that 60% of trials presented an ‘X’, 20% the ‘Q’ and 20% the ‘O’. Each block consisted of either the Go condition or No-Go condition. This task was designed to provide a boring task context commonly used in studies of ongoing experience.

##### 2.2.7.2. Self/Other adjective rating paradigm

This task was based on that used by de Caso and colleagues (de Caso, Karapanagiotidis, et al., 2017; de Caso, Poerio, Jefferies, & Smallwood, 2017), and is similar to other self-reference effect paradigms (Craik et al., 1999; Kelley et al., 2002; Vanderwal, Hunyadi, Grupe, Connors, & Schultz, 2008). Participants were instructed to attend to the centre of the screen, as they viewed adjectives, presented one word at a time. Each trial consisted of a probe signalling for the participant to either judge the following stimuli in accordance with the referent (self or other) or indicate whether the following stimuli was written capitalised. In experimental trials participants had to indicate whether they would associate the word presented with a referent or not. In one condition participants made judgements in relation to themselves (self condition) and another condition they made judgements in relation to a significant other (social cognition condition). Participants were verbally instructed to think of a single friend. In control trials participants indicated whether the words shown were written in uppercase or not. Each block consisted of participants making judgements about themselves (self condition) or friend (social cognition condition). The words used in this task paradigm were selected from a list of normalised personality trait adjectives with the highest meaningfulness ratings from Anderson and colleagues (Anderson, 1968), as used in de Caso and colleagues (de Caso, Karapanagiotidis, et al., 2017; de Caso, Poerio, et al., 2017). Each adjective list consisted of negative adjectives (50%) and positive adjectives (50%). Participants saw a different list of words in each block. This task was designed to engage participants in social cognition, making judgements in relation to themselves as well as a significant other.

##### 2.2.7.3 Semantic task paradigm

This task was adapted from the task paradigm used by Rice and Colleagues (Rice, Hoffman, Binney, & Lambon Ralph, 2018). Participants were instructed to attend to the centre of the screen, as they viewed four categories of stimuli: i) pictures of people, ii) pictures of places, iii) written people iiii) written places. The stimuli used in this task consisted of trials with an 85% or greater accuracy from Rice and colleagues (Rice, Hoffman, Binney & Lambon Ralph, 2018) as in Gonzalez Alam et al. (in review). Each block consisted of the two picture or word categories. Each trial consisted of a probe signalling to judge the following stimuli on being European or located high on the screen. In experimental trials participants had to indicate whether the stimuli shown were European. In control trials participants had to indicate whether the stimulus shown was located high on the screen (above the fixation cross). This task was designed to engage participants in making semantic judgements with stimuli of varying modality.

#### 2.2.8. Cambridge neuropsychological test automated battery

The Cambridge neuropsychological test automated battery (CANTAB), a computerised cognitive assessment and data collection tool, was used to collect measures of executive function, memory, emotion and social cognition in participants (CANTAB® [Cognitive assessment software]. Cambridge Cognition (2019). All rights reserved. www.cantab.com). Participants completed the Cambridge gambling, emotional recognition, intra-extra dimensional set-shift and spatial working memory tasks once during the task session using i-Pads. Full details of the tasks below can be found at www.cantab.com.

##### 2.2.8.1. Cambridge Gambling Task

In the Cambridge Gambling Task (CGT) participants were instructed to attend to the screen as they viewed 10 boxes, some which were red and others blue. The ratio of red to blue boxes varied on a trial basis. On every trial a token was hidden under one of the 10 presented boxes. A counter showing points was also presented on the screen. Participants started the task with 100 points. They had to guess whether the token would be hidden under a red or blue box using a forced choice key response (red or blue) while also betting points on their response. In the first block the points increased, in the second block the points decreased. If the answer was correct participants earned the points shown on the counter, if they were incorrect they lost the points shown. This task took ~ 18 minutes. The Cambridge gambling task was designed to measure decision-making as well as risk-taking behaviour. Due to a technical error half of the recruited participants (35 participants) completed a 6 minute longer version of this task with 36 more trials.

##### 2.2.8.2. Emotional recognition task

In the emotional recognition task (ERT) participants were presented with faces (computer-morphed images producing an average face composed from pictures of a range of individuals), which expressed one of the six basic emotions (anger, disgust, fear, happiness, sadness and surprise). They had to indicate which one of the 6 basic emotions the stimulus expressed using a 6-button forced choice response. This task took ~ 10 minutes. The emotional recognition task was designed to measure participants’ ability to identify each of the six basic emotions.

##### 2.2.8.3. Intra-extra dimensional set shift task

In the intra-extra dimensional set shift (IED) task participants were presented with two categories of stimuli (pink shapes and white lines). The start of the task consisted of simple stimuli which only varied in one category, for example, two white lines of different shapes. As the task progressed participants were presented with more complex stimuli, for example, stimuli consisted of a mixture of the two categories such as white lines superimposed on pink shapes. Participants had to use feedback (a high pitched tone indicating a correct answer or a low pitched tone indicating an incorrect answer) to figure out the rule used to identify the correct stimulus on each trial. The rule changed after 6 correct responses. At the start of the task, one dimension was the focus of the rule (e.g. pink shapes), and as the task progressed participants had to adapt to the change in focus of the rule (e.g. white lines become the focus). This task took ~ 7 minutes. The intra-extra dimensional set shift task was designed to measure participant ability to attend to a particular category of stimuli and later shift this attention to categories of stimuli that were ignored. Due to a technical error 5 participants reaction times were not recorded.

##### 2.2.8.4. Spatial working memory task

In the spatial working memory task (SWM) participants were presented with boxes. They were instructed to search in the boxes to identify a hidden token using the process of elimination. If a token was hidden under a box, it was not hidden under that same box in following trials. Participants were presented with an increasing number of boxes as the task progressed (4, 6 and 8 boxes). This task took ~ 4 minutes. The spatial working memory task was designed to measure search strategy and memory error (searching in a box that contained a token on a previous trial and searching in a box twice in the same trial).

### 2.3. Data analysis

#### 2.3.1 Principal component Analysis

Analysis of the MDES data was carried out in SPSS (Version 25, 2019). Principal Component Analysis (PCA) was applied to the scores from the 13 experience sampling questions (see Table 2) comprising the probes for each participant in each task. This was applied at the trial level in the same manner as in our prior studies (Konishi et al., 2017; Ruby, Smallwood, Engen, & Singer, 2013; Ruby, Smallwood, Sackur, & Singer, 2013; Smallwood et al., 2016; Turnbull, Wang, Schooler, et al., 2019). Specifically, we concatenated the responses of each participant for each trial into a single matrix and employed a PCA with varimax rotation. Components were selected based on the scree plot (Figure 1). All tasks were included to examine thought patterns across the range of task states measured. Due to technical issues, eight participants had seven MDES probes rather than eight in the affective TV-based task and one had three probes from the CANTAB task rather than four. Two participants’ CANTAB probes were excluded from analysis due to task incorrect completion of the MDES probes. Finally, due to technical issues two participants completed the sessions in a different order compared to the rest of the cohort.

**Figure 1.**
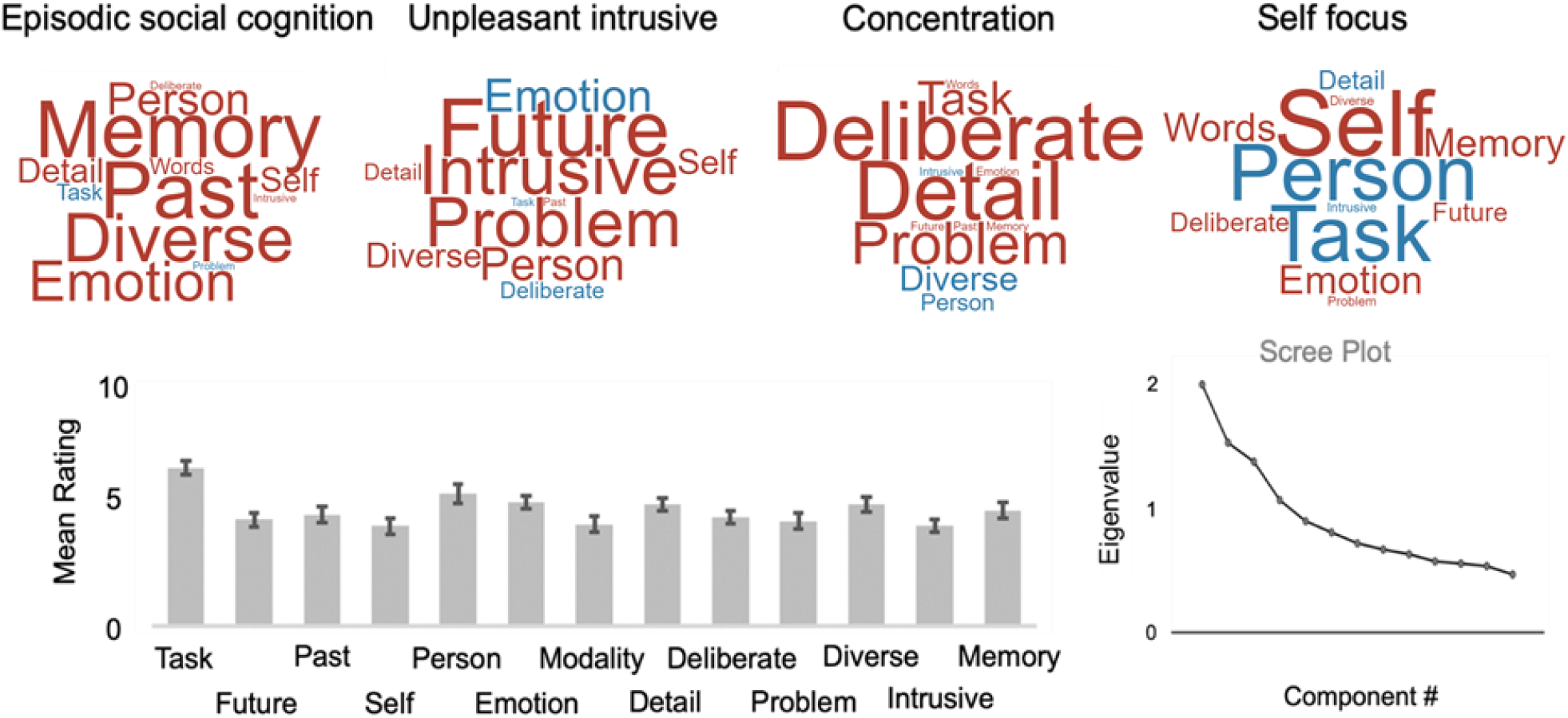
Decomposition of the experience sampling data collected in this study revealed four components across all conditions. Based on their loadings the four components were summarised as “Episodic Social Cognition”, “Unpleasant Intrusive”, “Concentration” and “Self Focus”. The word clouds in the upper panel summarise these loadings in which the colour of the word describes the direction of the relationship (red = positive, blue = negative) and the size of the item reflects the magnitude of the loading. The bar-plot in the lower panel shows the mean ratings for each item that these components are derived from. The scree plot for this decomposition is presented in the lower right panel. Error bars represent 99.6% CI.

#### 2.3.2 Linear Mixed Model

A linear mixed model (LMM) was implemented in SPSS version 25 to examine whether the dimensions of thought identified in the PCA varied significantly across the task conditions. In this analysis we performed four separate models in which each of the components identified in the PCA was an outcome measure, and the task conditions were included as conditions of interest. In these models we included probe number, order, and day of testing as nuisance co-variates of no interest. The participants’ intercept was treated as a random factor.

#### 2.3.3 Repeated Measures Analysis of Variance

A repeated measures analysis of variance (ANOVA) was run to examine how the contextual influences on each dimension of thought related to the measures of affective style recorded in the current study (depression, state and trait anxiety). The outcome variables were the mean scores for each participant for each PCA component (a total of 4 variables), and the explanatory variables were participants’ scores on the measures of affective style (depression, state and trait anxiety).

### 2.4. Data and code availability statement

Multidimensional Experience Sampling data is available upon request from the authors via email.

Ethical approval conditions do not permit public sharing of raw data as participants have not provided sufficient consent. Data and analysis scripts can be accessed by contacting the corresponding author, Delali Konu. Data will be accessible upon request within accordance with General Data Protection Regulation (GDPR).

## 3. Results

To provide a compact low dimensional representation of the experience sampling data we applied Principal Component Analysis (PCA; see Methods). Based on the scree plot we observed an elbow after four components, which in total accounted for 53.22% of the total variance. The loadings on these components are presented as word clouds in Figure 1. Component One accounted for 15.28% of the variance and reflects patterns of positively valenced episodic social cognition (“episodic social cognition”). Component Two accounted for 13.91% of the experience sampling data and reflected a pattern of negatively valenced intrusive thought (“unpleasant intrusive”). Component Three accounted for 12.31% of the variance and reflects high loadings on deliberate detailed task focus (“concentration”). Finally, Component Four accounted for 11.73% of the overall variance and reflected a pattern of off-task self-relevant cognition that has negative loadings on the “Person” feature but positive loadings on “Self” feature, separating thinking about the self from thinking about other people (“self focus”).

Having determined four dimensions of ongoing thought in the experience sampling data we next examined whether these varied significantly across the task environments. We addressed this question using a linear mixed model (LMM; see Methods). We performed four separate models in which each of the four components were an outcome measure.

This analysis revealed a significant influence of task condition on the distribution of each component (Component One, F (14, 2205.64) = 86.89, p = <.001; Component Two, F (14, 2205.54) = 27.39, p <.001; Component Three, F (14, 2205.72) = 37.70, p < .001 & Component Four, F (14, 2205.81) = 123.17, p <.001). The results of this analysis are presented in Figure 2, both in the form of a bar plot summarising the beta weights from the model (including confidence intervals), and in the form of word clouds. In each bar graph the conditions are ordered by their relative influence on the relevant dimension of thought. It can be seen from Figure 2, that Component One (episodic social cognition) was most common in the task which required participants to rate the applicability of items to a significant other (friend) and lowest weighting in more complex tasks (e.g. working memory), as well as in the affectively toned TV clips (action and suspense). Component Two (intrusive thought) was most prevalent in the affectively-toned TV clips. Component Three (concentration) was most prevalent in demanding tasks (working memory and switching) and least prevalent in the tasks with a narrative but without strong affective ties (audio and video documentary conditions). Finally, Component Four (self-focus) was prevalent during self-reference and gambling, sustained attention tasks, and was absent from the affectively toned TV clips (action and suspense).

**Figure 2.**
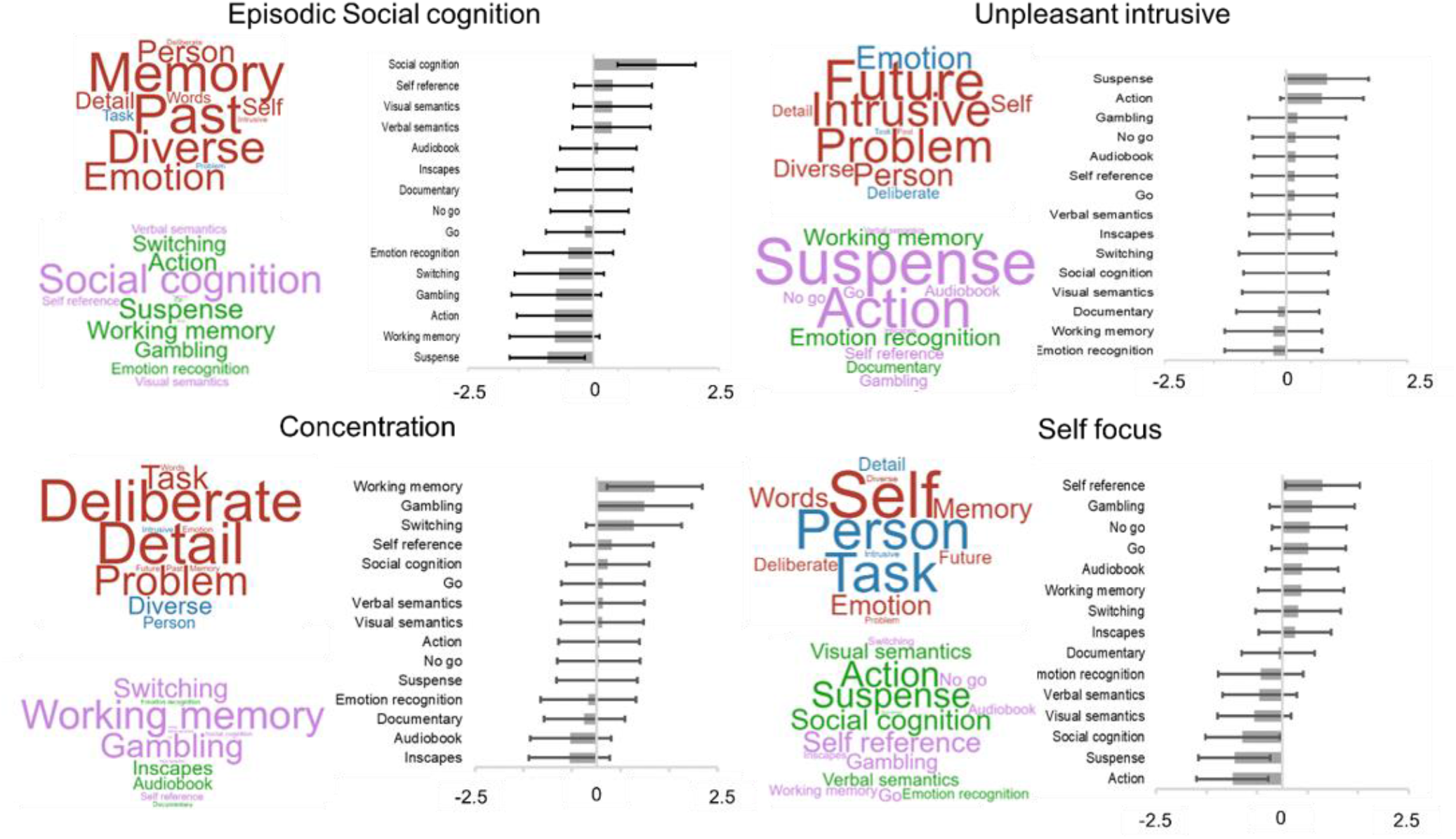
Results of a linear mixed model (LMM) examining the situational variance in the four patterns of thought identified using PCA. In each panel the top word cloud reiterates the loadings on each thought pattern for the convenience of the reader (the colour of the word describes the direction of the relationship; red = positive, blue = negative, and the size reflects the magnitude of the loading). The lower word cloud highlights the loadings of this pattern in each task as described by the parameter estimates from the LMM (the colour of the word describes the direction of the relationship; purple = positive, green = negative, and the size reflects the magnitude of the loading). The bar plot shows the same data and reports the confidence intervals for these estimates (p < .05, corrected for family wise error). Error bars, therefore, represent 99.7% CI. Action = action (affective TV-based), Audiobook = audio (documentary TV-based), Documentary = documentary (documentary TV-based), Emotional Recognition = ERT (CANTAB), Gambling = CGT (CANTAB), Go = go (Go/No-go), Inscapes = Inscapes (documentary TV-based), No-go (Go/No-go), Self reference = self (self paradigm), social cognition = social cognition (self paradigm), suspense = suspense (affective TV-based), Switching = IED (CANTAB), Verbal semantics = word (Semantic paradigm), Visual semantics = picture (Semantic paradigm), Working memory = SWM (CANTAB).

**Figure 3.**
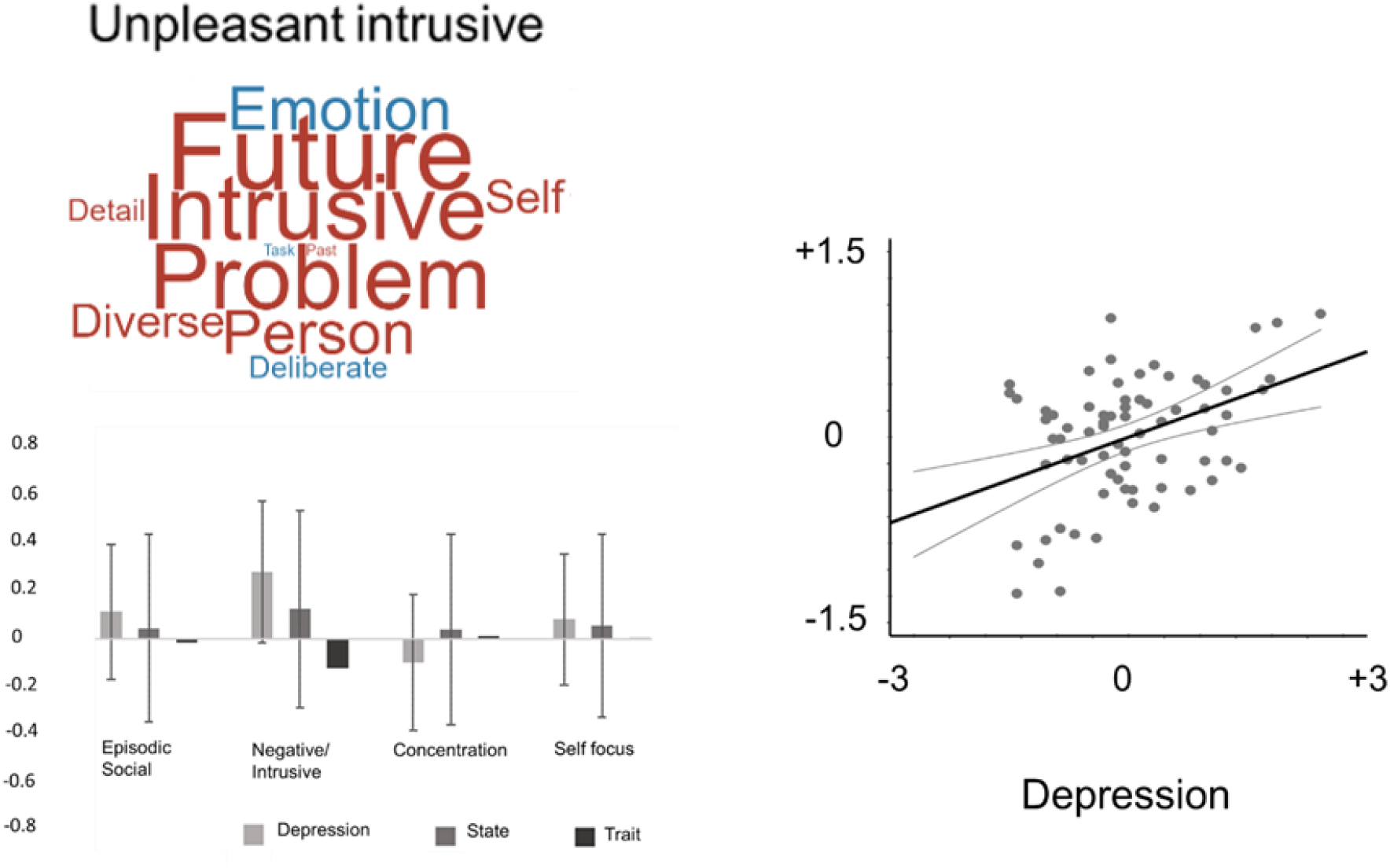
The association between patterns of thought and measures of affective disturbance (depression, state and trait anxiety). The bar graph summarises the beta weights from the model describing the average contribution of depression, state and trait anxiety as described by its parameter estimate and associated confidence intervals. We found that patterns of unpleasant intrusive thoughts were positively associated with levels of higher depression (p<.05). The scatterplot shows the distribution of this relationship in which each point is a participant. Error bars represent 95% CI.

Having determined the contextual influences on each pattern of thought we next examined how they related to the measures of affective style recorded in our experiment (depression, state and trait anxiety). This revealed a significant Component by Depression interaction (F (3,198) = 2.95, p<.05). Further analysis indicated that higher levels of depression were associated with higher scores on the intrusive thought component (r =.418, p<.001).

## 4. Discussion

Our study set out to understand how thought patterns vary across a wide range of task environments, including those which encompass both simple and complex laboratory tasks, as well as more realistic everyday task situations such as TV-based conditions with varying affective components. We used MDES to characterise patterns of thought during blocks of task performance along multiple dimensions (see Table One). We applied Principal Components Analyses (PCA) to these data to identify the latent dimensions that best described these variables. We found that this approach revealed four dimensions that we summarised as “episodic social cognition”, “intrusive negative thought”, “detailed deliberate thought” and “self focus“. Three of these dimensions, “episodic social cognition”, “detail and deliberate” and “intrusive negative”, were similar to ones observed in our prior study using the same set of questions and assessing experience in a simple signal detection paradigm (Konu et al., 2020). In the current study, each of the four dimensions captured by PCA varied across the task environments we studied. The pattern of episodic social cognition was most evident when participants think about features of a significant other (their friend) and least prevalent while watching affective TV clips and during working memory tasks. Complex demanding tasks (working memory, switching or gambling) were linked to patterns of detailed deliberate thoughts, replicating a pattern seen in our prior studies in which we found that thoughts had this property with increasing working memory demands (Sormaz et al., 2018; Turnbull, Wang, Murphy, et al., 2019). Unpleasant intrusive thoughts were most common while watching TV clips with affective features. Finally, when participants engaged in self-referential thought patterns, they reported thinking most about themselves. This thought pattern was least important while watching affectively toned TV clips or thinking about other people. We also found that patterns of intrusive thoughts were more prevalent in the self-reports of individuals with higher levels of depression. These data therefore show that thought patterns can reflect the influence of both testing conditions and individual affective style which can both be captured using a low dimensional space described by PCA. We consider these data in the context of prior work examining the features of different thought patterns.

First, these data add to a growing body of evidence that patterns of ongoing thought are heterogeneous across individuals and different environments. Our PCA analysis highlighted four different thought patterns and each encompass features that mind-wandering could be argued to have – stimulus independent features (loadings of memory in Component 1), intrusive features (high loadings on Component 2), the absence of a deliberate assessment of task-relevant information (i.e. Component 3) and a trade-off between task focus in favour of self-relevant sources of information (Component 4). Importantly, we found evidence that each of these different experiential features varied in their prominence across the task conditions. These results suggest that there may be multiple different patterns of experience which may be distributed in a complex way across different task contexts. Work examining the phenomena of mind-wandering has reached a conceptual impasse since there is no consensus on defining features of the experience, or whether these are even necessary (Christoff et al., 2018; Paul Seli et al., 2018). In this context, our application of PCA to experience sampling data from across a wide range of task environments may provide a helpful way to empirically unpack the complexity and richness of the state space that experience sampling allows experimenters to examine. Moving forward, therefore, our study adds to a growing call for both conceptual and definitional clarity when using experience sampling to define experiential states, and to characterise their features or associated underlying thought processes.

Second, it is possible that the heterogeneous space identified in our analysis partly explains why patterns of ongoing thought findings map with varying degree from the laboratory to daily life (Ho et al., 2020; Linz et al., 2019; McVay, Kane, & Kwapil, 2009). We found that different patterns of experience tended to be expressed in a complex manner across task environments. For example, patterns of off-task self-relevant thought were common in tasks that emphasise the self as a target (self-reference) or indirectly (gambling or boring sustained attention tasks) relative to when people watched extracts of affectively engaging TV clips. In contrast, detailed task focus was highest in complex tasks (working memory) and lowest in TV clips and audiobooks with fewer affective features. In laboratory studies of “mind-wandering”, researchers employ sustained attention or working memory tasks, while in daily life it is likely that listening to audiobooks or watching TV clips is a more common activity. Based on our data, systematic variation in the tasks used in cognitive experiments from those that participants tend to engage in their day-to-day lives may be one important factor to consider when trying to map between the laboratory to the real world. Importantly we have recently used PCA to map similarities and differences between patterns of ongoing thoughts recorded via experience sampling in the lab and in the real world (Ho et al., 2020). In the future it could be possible to use techniques like PCA to identify task environments which best capture the patterns of thoughts that people encounter in daily life and use these in the laboratory to gain a more ecological perspective on cognition in the real world (Matusz, Dikker, Huth, & Perrodin, 2019; Smilek, Eastwood, Reynolds, & Kingstone, 2007).

One specific implication of our result concerns the choice of task to use in the laboratory to study mind-wandering. One tacit assumption within the literature is to use tasks that lack compelling task demands (i.e. the Go and No-Go tasks or relatively dry narratives such as the documentaries) because they may promote off-task states. Consistent with this, we found that these tasks did not promote a state of task focus as effectively as did the working memory tasks (See Figure 2). Instead, these contexts emphasised patterns of self-focus to a much greater degree than did affectively toned TV-clips. These observations support the consensus within the literature that these paradigms provide a fertile context in which to study experiences such as mind-wandering. Importantly, however, our study qualifies the assumption that although the relative absence of patterns of concentration in these non-demanding tasks suggests that they provide paradigms that are well suited for understanding the generation of personally relevant thought, they may not be best suited to understanding how individuals maintain states of concentrated task performance. It is possible that this is why motivation plays such an important role in states of mind-wandering in tasks like the SART (P. Seli, Cheyne, Xu, Purdon, & Smilek, 2015).

Third, our study adds to a growing body of evidence highlighting that there may be multiple mechanisms through which task conditions can influence ongoing thought. Prior studies show that patterns of social-episodic thoughts are reduced when individuals engage in complex external tasks, in which context experience is dominated by a pattern of detailed task focus (Turnbull, Wang, Murphy, et al., 2019; Turnbull, Wang, Schooler, et al., 2019). Our study is broadly consistent with these prior findings since we find a similar component which is most prevalent when participants engage in socially motivated tasks, and is minimal in both complex tasks (working memory) as well as in affectively toned TV clips. This pattern of suppression of self-generated material in conditions of higher task demands is usually interpreted in terms of the need to maintain task focus to perform more complex tasks (Teasdale et al., 1995; Teasdale, Proctor, Lloyd, & Baddeley, 1993). In contrast, during reading, difficult texts can be hard to maintain focus on (Feng, D’Mello, & Graesser, 2013), while interesting texts may be easier to attend to (Smallwood et al., 2009; Unsworth & McMillan, 2013). Notably patterns of self-focused experience exhibited a different contextual distribution – these were more common in working memory tasks than in affectively toned TV clips (See Figure 2). It is possible that this difference relates to the role that salience plays in capturing ongoing thought. It has been suggested that individuals often focus on their current concerns when they escape the here and now (Cox & Klinger, 2004; Klinger, 1987). We speculate that it may be the saliency of the information in the affective TV clips which helps individuals to anchor attention in the here and now and briefly escape from their own worries, a perspective that is supported by the fact that this same pattern of thought was generally elevated in less happy individuals for whom current concerns may be relatively high (Ruehlman, 1985).

## 5. Conclusion and Limitations

Although our study establishes the role that both individuals and situations play in patterns of ongoing thought, it leaves several important questions unanswered. First, our study was composed of university educated students and this limits the degree to which these results would generalise to older populations for whom patterns of thoughts are known to be different (Giambra & Grodsky, 1989). Second, although our design allowed us to understand the influence of both task environments and an individuals’ affective style on patterns of ongoing thought, it remains unclear how these two processes interact. It would be useful in the future, for example, to understand whether the association between depression and patterns of unpleasant thought is stronger or weaker in the presence or absence of threat in the environment. Third, for pragmatic reasons the number of measures of experience in each task was uneven. Although our analysis suggests that we had sufficient power to discriminate the pattern of thoughts across different situations, it remains possible that the amount of variance captured by each PCA could be influenced by the number of samples in each context, and it is also possible that this may also influence qualities of the patterns themselves. Fourth, although the use of TV-clips as a task environment enables us to test for patterns of thought in a more naturalistic setting, there is still room to develop more naturalistic task environments. In this regard, it is important to note that our selection of the questions to assess ongoing thought and the tasks we used or the measures of affective style is in no way a comprehensive description of either the types of tasks people perform, or the thoughts they have. It is likely that there are many task environments that our study does not capture, many aspects of experience that our questions did not query and multiple features of an individuals’ disposition that influence their experiences. However, our study uses a greater range of task environments and experience sampling questions than is standard in this type of work and thus highlights that the study of limited aspects of experience in only a subset of possible task environments are likely to prohibit our ability to fully appreciate the different patterns of ongoing thoughts that individuals can have. Thus, our study highlights the need to broaden the tasks we use to study ongoing thought and raises the need to develop a conceptual framework within which the scientific study of self-generated thought can be embedded.

## CRediT authorship contribution statement

Delali Konu: Conceptualization, Data curation, Formal analysis, Investigation, Methodology, Project administration, Visualization, Writing – original draft. Brontë Mckeown: Conceptualization, Data curation, Formal analysis, Investigation, Methodology, Project administration, Software, Writing – original draft. Adam Turnbull: Conceptualization, Data curation, Formal analysis, Investigation, Methodology, Project administration, Software, Writing – original draft. Nerissa Siu Ping Ho: Software. Tamara Vanderwal: Conceptualization, Writing – original draft. Cade McCall: Conceptualization, Methodology, Project administration, Supervision, Visualization, Writing – original draft. Steven P. Tipper: Conceptualization, Writing – original draft. Elizabeth Jefferies: Conceptualization, Writing – original draft. Jonathan Smallwood: Conceptualization, Formal analysis, Funding acquisition, Methodology, Project administration, Supervision, Visualization, Writing – original draft.

## Notes

This work was supported by European Research Council awarded to JS (WANDERINGMINDS – 646927).

### Competing Interest Statement

The authors have declared no competing interest.

